# Fine root endophytes forming winter mycorrhiza

**DOI:** 10.1101/2025.01.10.632310

**Authors:** Takeshi Sentoku, Yuki Komatsuda, Hayato Shimada, Michio Arai, Yoshihiro Kobae

**Affiliations:** Arai Helmet Ltd., 2-12, Azuma-cho, Omiya-ku, Saitama-shi, Saitama 330-0841, Japan; Department of Sustainable Agriculture, Rakuno Gakuen University, 582, Bunkyodai Midori-cho, Ebetsu, Hokkaido 069-0836, Japan; Agrisystem Ltd., 15-8 Higashi-Memuro, Memuro-cho, Kasai-gun, Hokkaido, 082-0005 Japan

## Abstract

Arbuscular mycorrhizal fungi, classified in the subphylum Glomeromycotina, are obligate symbionts that depend on photosynthetic products from plants. There is substantial evidence that AMF support plant and crop growth in natural and agricultural ecosystems. Fine root endophytes (FRE) also co-occur in plant roots with AMF in all but tropical environments. However, their presence remains poorly recognised, and their lifestyle and functionality remain largely unknown. Our analysis demonstrates that, in contrast to AMF, FRE colonise plants during the winter season, when photosynthesis is more challenging. It is also noteworthy that FRE was unable to colonise the roots of spring-sown crops. The results of our laboratory pot experiments demonstrated that low temperatures are not sufficient for FRE colonisation. Furthermore, we observed that FRE, in contrast to AMF, exhibited increased colonisation in soils containing metabolically inactive plants. Our results suggest that FRE does not contribute to the increase of ecosystem biomass through direct photosynthesis in summer, but may play an overlooked role in the formation of mycorrhiza-based soil ecosystems in winter. However, this winter ecosystem can be disrupted by conventional bare fallow management of the field, interrupting the annual cycle.

## Introduction

Arbuscular mycorrhizal fungi of the subphylum Glomeromycotina colonise plants in a wide range of environments, from warm to cold climates and from poor to fertile soils, and their dynamics and functions in natural and agricultural ecosystems are well documented (Douds and Millmer, 1999; Powell and Rillig, 2018). AMF forms arbuscules in the root cortical cell of a wide range of plant lineages, from liverworts to angiosperms, and delivers nutrients (particularly phosphate) to the plants (Karandashov and Bucher, 2005). In turn, the photosynthetic products derived from plants contribute to mycorrhizal development and the accumulation of organic matter in the soil (Frey, 2019). On the other hand, knowledge about the ecology and function of fine root endophytes (FRE) remains limited (Orhcard et al., 2017b; Walker et al., 2018). Reasons for the delay in this research include: i) mycelia of FRE in plants are thinner than those of AMF and therefore difficult to observe, ii) except in some “extreme” environments, the roots of most plants are generally dominated by AMF colonisation (Orchard et al., 2017a), requiring the identification of FRE within AMF background needs high power microscope and experience of observation, iii) the morphological characteristics of FRE mycelia are not systematically defined, iv) after sampling, FRE colonisation in roots decrease significantly compared to AMF (Orchard et al., 2017c), v) FRE spores are very small (< 20 µm) compared to AMF (30 µm – 1 mm) and it is impossible to separate FRE from soil by standard methods (Thippayarugs et al., 1999), making it difficult to isolate and culture FRE (Orchard et al., 2017b), and vi) genomic and phylogenetic information on FRE is insufficient (Seeliger et al., 2024), making difficult to deepen the understanding the specific characteristics of FRE (Lutz et al., 2025). Despite these difficulties, years of careful observation and efforts to isolate and culture FRE have only recently begun to report its biologically unique and interesting features (Bonfante and Venice, 2020; Prout et al., 2024).

## FRE mycorrhiza increases in winter

FRE tends to be observed in soil environments with lower temperatures (Orchard et al., 2017b), waterlogging (Orchard et al., 2016), lower available soil phosphate (Albornoz et al., 2021) and acidic soil condition (Arines et al., 1988). Kowal et al. (2020) studied FRE colonising a fern species (*Lycopodiella inundata*) from spring to autumn in heathlands in England and the Netherlands, and reported that FRE was more common in autumn when temperatures were low and precipitation was high. Albornoz et al. (2021) studied AMF colonisation of clover (*Trifolium subterraneum*) throughout Australia in winter and reported that FRE is more likely to be observed in temperate agricultural areas, but not in the tropics. However, the AMF that first colonise the roots of seedlings in Japanese temperate agricultural soils are often normal AMF, not FRE. Kowal et al. (2020) pointed out the importance of studying the seasonal dynamics of FRE and the risks of analysing only temporal points, but to our knowledge there is no report of fixed-point analysis of FRE over one year. Therefore, we investigated the proportion of AMF and FRE colonising the roots of different plant species over time for two years, from January 2023 to January 2025, in the field in Saitama, Japan, where FRE colonisation of plant roots is observed in autumn. This field has a diverse vegetation as a cover crop as well as crops. While there were differences in temperature and precipitation from year to year, there was a clear trend that FRE was rarely observed in summer and frequently observed in winter (**Figure 1**).

**Figure 1.**
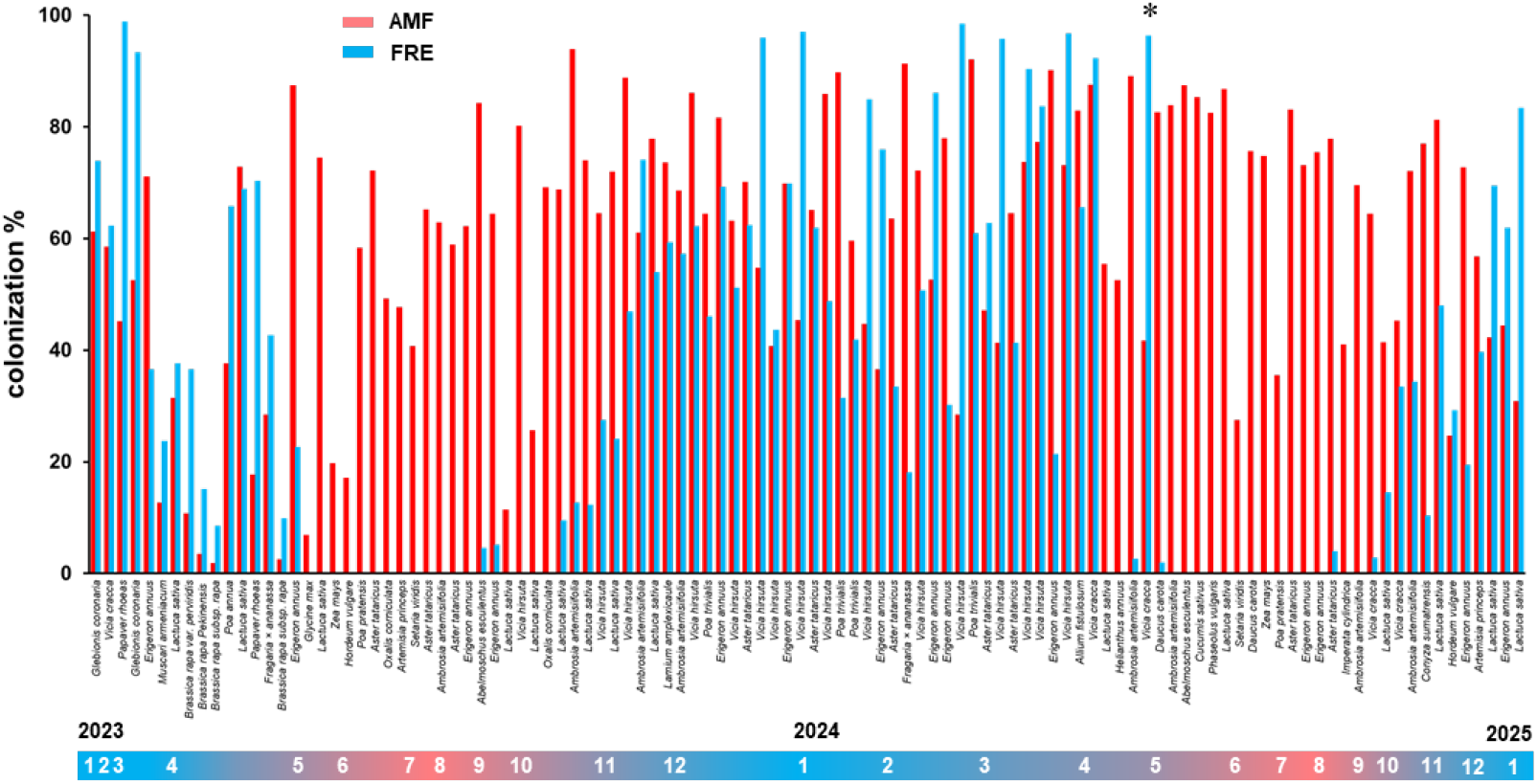
Change in colonisation rates of AMF and FRE during two years. A total of 28 plant species were studied over a two-year period from January 2023 to January 2025 in agricultural field in Saitama, Japan. Mycorrhizal fungi colonising the roots were stained with trypan blue and their percent root length colonisation were assessed by modified gridline intersection method. The roots of all 21 plant species observed in winter season were co-colonised with arbuscular mycorrhizal fungi (AMF) and fine root endophytes (FRE). Plant indicated by asterisk retained overwintering mycorrhizal roots colonised with FRE, while FRE was rarely found on the roots of plants in summer season.

FRE and AMF co-colonise the roots of all 21 plant species examined in winter (Asteraceae, Fabaceae, Papaveraceae, Liliaceae, Brassicaceae, Poaceae, Malvaceae, Apiaceae, and Lamiaceae). In May, AMF predominantly colonised the roots of seedlings sown, while FRE colonisation was rare. The rare roots that were well colonised with FRE were the roots of overwintered plants. FRE did not show high colonisation rates in the roots of summer plants until autumn. In winter, although the trypan blue staining of coarse intraradical hyphae (> 3 µm in diameter) of AMF (Crush, 1975) was weaker (**Supplementary Figure 1a**), they were certainly found in the roots and the colonisation level was high in many plants. Many of the arbuscules derived from the coarse intraradical hyphae appeared to have collapsed or stopped branching: i.e. the proportion of these morphologies appeared to be increasing: this suggests that the cycle of intracellular colonisation of AMF is retarded (Pearson et al., 1991). In contrast, FRE was very active in colonising roots in winter, forming mycelial structures typical of FRE such as fan- or comb-like structures, and branching thin hyphae (< 1.5 µm in diameter) which often form small vesicles (5–10 µm in diameter) (Thippayarugs et al., 1999; Orchard et al., 2017b; Walker et al. 2018) (**Supplementary Figure 1b**).

## FRE colonisation is not induced by low temperature alone

To confirm the field observations in a laboratory environment, red clover was grown in 20 ml pots filled with soil from the same field (passed through a 2 mm sieve) under two conditions: high temperature (26°C with light for 16 h / 23°C in darkness) and low temperature (18°C with light for 10 h / 14°C in darkness) to determine the type of AMF that colonise the roots. Under the high temperature condition, AMF colonisation (hyphopodium or infection unit) was observed on the fourth day of cultivation and the number of colonisation increased during the following four days, while FRE colonisation was rarely observed (**Figure 2a**). Under low temperature conditions, the onset of AMF colonisation was delayed by about 3 days compared to high temperature, but the number of colonisation increased during the following 4 days (**Figure 2b**). FRE showed fewer colonisation, but their number did not increase during the following 4 days. To determine whether prolonged low-temperature cultivation promotes FRE colonisation, the same pots were grown for 30 days under three conditions: 25°C with light for 16 h, 18°C with light for 10 h, and 8°C with light for 10 h. Cultivation at the lowest 8°C, significantly reduced AMF colonisation, and FRE colonisation was also reduced and not promoted at lower temperatures **(Figure 2c**). A 30-day cultivation of red clover in small 20 ml pots (**Figure 2c**) showed yellowing leaves, nutrient deficiency or senescence. Prolonged cultivation in small pots can stress the plant, severely limiting root elongation and depleting some of the soil nutrients. Interestingly, red clover showed a relatively high colonisation of 10% under these stress conditions. Therefore, we predicted that the host plants with reduced metabolic activity would be colonised by FRE. However, the fact that FRE colonises seedlings sown in autumn in agricultural fields (**Figure 1**) suggests that if the infectivity of FRE in the soil in autumn is enhanced by the presence of senescent plants, FRE can colonise nearby plants regardless of their physiological state. To assess this possibility, we first cultivated red clover in 20 mL pots and sowed lettuce under conditions where the metabolic activity of red clover gradually decreased, and examined the AMF colonising red clover and lettuce roots (**Figure 2d**). Red clover roots after 4 weeks of cultivation were mainly colonised with AMF, but the percentage of roots colonised with AMF gradually decreased after 8 weeks of cultivation, and the percentage of roots colonised with FRE increased (**Figure 2e**). Roots of 2-week-old lettuce seedlings mixed with 4-week-old red clover were predominantly colonised with AMF, as was the case with red clover, while roots of 2-week-old lettuce seedlings mixed with 8-week-old red clover were similarly colonised with AMF and FRE (**Figure 2f**). This pot-culture experiments confirmed that AMF and FRE lifestyles are quite different in the laboratory environment as well as in the agricultural field.

**Figure 2.**
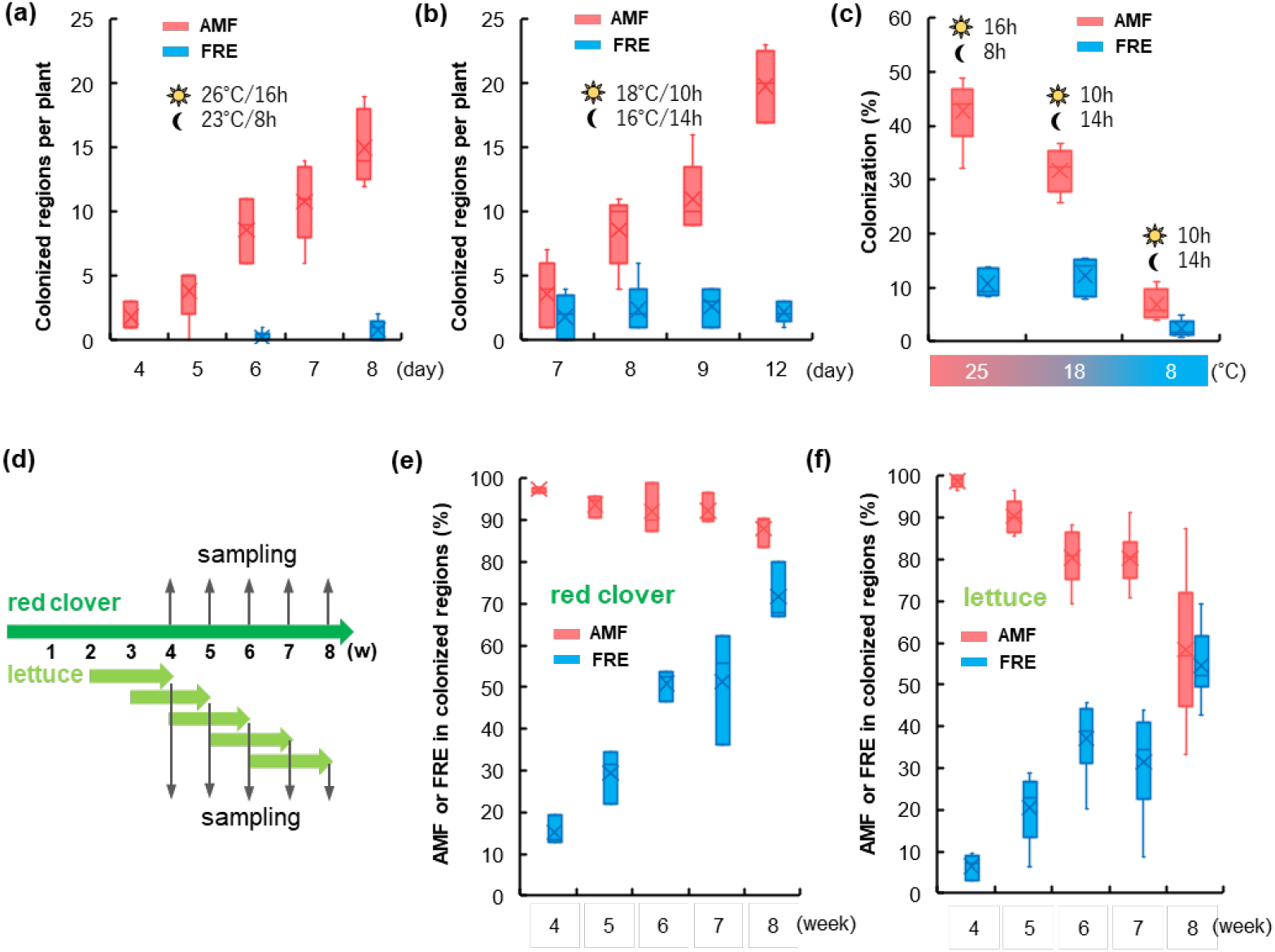
FRE colonisation in pot culture. Red clover was grown in pots under different environments to investigate the conditions under which fine root endophytes (FRE) can colonise the plants. The 20 mL pots were filled with soil from Saitama (Japan) agricultural field after passing through a 2 mm sieve. **(a)** Red clover was grown at high temperature (16 h Light at 26°C/8 h Dark at 23°C) and sampled after 4, 5, 6, 7 and 8 days to count the number of colonised region of arbuscular mycorrhiza fungi (AMF) or FRE per plant (n=6). **(b)** Red clover was grown at low temperature (10 h light, 18°C/14 h dark, 16°C) and sampled after 7, 8, 9 and 12 days to count the number of colonised region of AMF or FRE per plant (n=6). **(c)** Prolonged low temperature-cultivation does not promote FRE colonisation. Red clover was grown at 25° C with 16 h light, 18°C with 10 h light and 8°C with 10 h light for 30 days. Percent root length colonisation of AMF or FRE was determined. **(d)** Design of the mixed culture of red clover and lettuce. Red clover was grown first in 20 mL pots and lettuce was sown beside red clover. **(e)** Red clover was sampled at 4, 5, 6, 7 and 8 weeks after sowing and the percentage of AMF and FRE observed at the colonised regions in the roots of one individual red clover was determined (n=3). **(f)** Lettuce was sown beside red clover in weeks 2, 3, 4, 5 and 6 of cultivation and grown for 2 weeks. The percentage of AMF and FRE observed at the colonised regions in the roots of one individual lettuce was determined (n=4).

## Future perspectives

The current study has identified winter active lifestyles of FRE through a year-round analysis. This study only looked at AMF in one area over time, so this result may have the geographical bias of a specific location. However, it does allow a detailed study of changes over time. The results show that FRE has completely different levels of colonisation in summer and winter. Although no conclusions about the geographical generality of the FRE lifestyle can be drawn from this study alone, previous reports have pointed out that FRE colonisation is often found microscopically and genetically in lower temperature environments (Tederssoo et al., 2014; Orchard et al., 2016; Albornoz et al., 2020;), and our surveys of more than 10 farmers in Japan have also shown that FRE is only found from autumn to spring.

The question raised by the fact that FRE produces winter mycorrhiza is how FRE obtains energy and nutrients to grow during the winter season when metabolic activity in the environment is decreased. Although it is not appropriate to extrapolate solely from the seasonal difference in AMF and FRE, we favour the hypothesis that FRE has a saprotrophic lifestyle, as suggested by some papers (Cole et al., 2024; Howard et al., 2024). Our pot experiments have shown that FRE can progressively colonise roots in soil environments containing plants with reduced metabolic activity. Since low temperatures alone do not promote FRE colonisation, other factors such as plant growth stage, available soil nutrients and the plant’s nutrient deficiency response may trigger the increase of FRE colonisation. We have not been successful in tracking where FRE are from spring to autumn. It is likely that most of the summer roots are new roots that have been supplied with photosynthates and have emerged, while last year’s overwintered roots have been degraded and released into the soil (Ruess et al., 2003). Decayed roots colonised with FRE, like AMF, can serve as inoculants in the autumn (Klironomos and Hart, 2002), or can be eaten by animals such as earthworms and insects, or degraded by decomposing microorganisms in summer (Gange, 2007). As a result, the intraradical mycelia and very small spores/vesicles of FRE released into bulk soil may have different dynamics in soil than AMF. FRE may be dormant during the summer, or it may live a plant-independent decomposing life. Understanding the summer lifestyle of FRE is also necessary to understand how FRE colonise plants in autumn.

Although certain FRE line (*Lyc-1*) has been shown experimentally to provide significant nitrogen to plants in axenic system (Hoysted et al., 2022), it is not clear how diverse the functions of FRE are in natural and agroecological systems (Jeffery et al., 2018). The diverse functionality of soil microorganisms is demonstrated by the diversity of their species (Lutz et al., 2025). However, knowledge of the species classification of FRE and the number of species is extremely limited. At least five morphological species have been recognised in FRE (Thippayarugs et al., 1999), but there seems to be no consensus on how to classify them morphologically (Seeliger et al., 2024). We also observed FREs with distinctly different morphologies throughout the year, but we could not find any trend as to whether this was due to species differences or whether different plant growth stages or seasons that caused the morphological changes. Although some FREs are classified as the subphylum Mucoromycotina (Prout et al., 2024) and called M-AMF, it is not certain whether all FREs observed in this study are classified as Mucoromycotina or whether they form arbuscules. Therefore, the term FRE is used throughout this paper.

Of the various FREs in the root, the symbiotic and functional fungal entities need to be identified. To this end, it will be important to find *in situ* molecular markers and specific staining methods that reflect AMF functionality (Kobae et al., 2010; Ivanov and Harrison, 2014). To obtain such functional markers, transcriptome analysis of FRE mycorrhiza may be useful, as has been done in the past for AMF studies. However, since FRE is often co-occurring with AMF, the functional information of AMF needs to be subtracted from that of FRE and AMF. Our pot culture system, which is transitioning from AMF only to AMF and FRE mix, is expected to help identify the function of FRE and in the future will lead to an understanding of the functional complementarity (Field et al., 2019) and nutritional interaction (Cole et al., 2024) underlying the co-occurrence of AMF and FRE.

Finally, the significance of the discovery of winter mycorrhiza is that it provides an opportunity to review conventional crop management practices, including tillage and excessive weeding (Turner and Meyer, 1991). Soil conservation farming have attracted attention in recent years as a deterrent to environmental change and global warming (Altieri et al., 2015). One of the most effective way to increase soil biomass, store organic matter, reduce greenhouse gas emissions and improve soil ecosystem services is to avoid leaving the soil bare, i.e. to cover it with green (plants) throughout the year (Abdalla et al., 2019), including in winter when crops are not cultivated. In commercially-oriented agriculture, priority has been given to the operational efficiency of cultivation, with little emphasis on the biological role of plants other than the crop to be harvested. Future research into the role of FRE could make a significant contribution to global environmental regeneration through the elucidation of their seasonal role in plant growth-supporting functions and in natural and agricultural soil ecosystems.

## Materials and methods

### Field plant and soil sampling

Field sampling was conducted from January 2023 to January 2025 at the cultivation field (35°54’23.4 ‘N 139°41’11.7 ‘E) of Alai Helmet Co Ltd, Omiya, Saitama, Japan. The soil type of the plot was homogeneous volcanic ash soil with a soil pH 6.2, available phosphate (Truog-P) 0 mg/100 g (below detection limit), exchangeable potassium 139 mg/100 g, nitrate nitrogen 0.48 mg/100 g, ammonium nitrogen 0.35 mg/100 g, exchangeable magnesium 33 mg/100 g, exchangeable calcium 222 mg/100 g and organic carbon 4.3 % (analysed in January 2023). Crops were randomly cultivated each year under no-tillage, with wild plants growing in the absence of crops. Crops and cover plants were sampled at different times of the year. The growth stages of the plants sampled varied from seedlings to overwintered plants. Their nutritional and growth conditions also varied. Collected roots were removed from the soil on site easily, transported to the laboratory within 30 minutes while maintaining humidity. Attached soil was carefully removed with tap water. Care was taken not to damage the roots. The roots were cut into 1 cm lengths with scissors and immersed in 50% ethanol for 3-16 hours.

### Pot experiments

The collected soil was passed through a 2 mm stainless steel sieve after one week of transport and storage at room temperature. It was then stored in a refrigerator for 6 months. Preliminary tests carried out during this 6-month period showed fluctuations in the colonisation activity of AMF, but no significant changes in the colonisation patterns of AMF and FRE or their colonisation proportions. A 25 ml self-standing clear polypropylene tube with a 5 mm hole drilled in the bottom was filled with 5 g of Akadama soil (tuff loam) (Setogahara Kaen, Gunma, Japan) as bottom soil to improve drainage, water retention and aeration. Soil stored for six months was added on the bottom soil. The soil contains high levels of coarse organic matter, which increases the volume of the soil at moderate moisture levels, but shrinks and reduces the volume when the soil absorb water. A tube of soil was therefore dropped twice onto a desk from a height of about 2 cm to compact the soil. The reduced amount of soil was then added. The process was repeated: the pot was placed in a container filled with 2 cm deep water in depth for 10 minutes to allow the pot soil to absorb the water.

Three red clover seeds were sown in the centre of the tube, covered with a small amount of calcined attapulgite clay to promote water absorption and stable germination of seeds, covered with a film to retain moisture and grown in a growth chamber with photosynthetic photon flux density (PPFD) of 100-150 µmol m^−2^ s^−1^, and varying temperature and photoperiod for each treatment. After 5 days, the film was removed, the plants were thinned to one plant and allowed to continue growing. In the mixed red clover and lettuce experiment, two lettuce seeds were sown next to 8 mm of red clover, covered with a small amount of calcined attapulgite clay to allow light to reach the seeds, covered with film to retain moisture, removed the film after 3 days, thinned to a single plant and continued growing. During the growing period, the pots were placed in a container with 2 cm deep water in depth for 10 minutes to allow the soil to absorb water.

### Trypan blue staining of AMF

Ethanol was replaced by 10% KOH, and the roots were heated in a water bath set at 95°C for a period of 10 minutes. The 10% KOH was then replaced with 2% HCl and left for a further 10 minutes at room temperature in order to acidify the interior of the roots. The roots were then treated with a solution of trypan blue (0.05% (w/v) trypan blue, lactic acid) and heated in a water bath at 95°C for 10 minutes. The roots were then rinsed with deionised water in order to remove any residual trypan blue, after which they were immersed in a lactoglycerol solution (consisting of 80% (v/v) lactic acid, 10% (v/v) glycerol and 10% (v/v) deionised water).

### Microscopic assessment of AMF colonisation

AMF colonisation was determined according to the method described by Giovannetti and Mosse (1980), unless otherwise stated. First, 20 roots were arranged in a parallel configuration on glass slides using forceps and a drop of lactoglycerol was added, followed by the placement of a coverslip. The specimens were observed at magnifications ranging from 200x to 1000x. The percentage of >100 intersections between the root and the grid that were colonised with AMF or FRE was calculated. The presence of AMF was evaluated through a comprehensive assessment of the morphology of the hyphopodium, the thickness of the intraradical mycelium (> 3 µm diameter), the presence of coils of exodermis and the linear extension of the mycelium in the cortical layer. FRE colonisation was evaluated based on the morphological description of many previous studies (e.g., Orchard et al., 2017b). The presence of FRE was evaluated through a comprehensive assessment of the morphology of the hyphopodium, the presence of fine intraradical hyphae with a diameter of < 2 µm, intercalary vesicles, intercellular comb-like structures that spread thinly across the cortex, intraradical hyphae that spread in a waterfall-like pattern from the hyphopodia, and the morphology of the mycelium that have minute branches and curves in the cortex. When it is difficult to decide whether it’s AMF or FRE, the intersection was classified as AMF x 0.5 and FRE x 0.5.

## Acknowledgements

This work was supported by JST-ALCA-Next Japan Grant Number JPMJAN24D3. The author would like to thank the many farmers who gave us the opportunity to carry out soil surveys on their land. The authors are grateful to the editor and to the anonymous reviewers for their constructive comments on the article.

## Author contributions

TS, MA and YK(Kobae) conceived and designed the investigation. TS, YK(Komatsuda) and HS undertook the experiments and analysed the results. ST and YK led the writing; all authors discussed results and commented on the manuscript. YK(Kobae) agrees to serve as the author responsible for contact and ensure communication.

## References

1 Abdalla, M., Hastings, A., Cheng, K., Yue, Q., Chadwick, D., Espenberg, M., … & Smith, P. (2019). A critical review of the impacts of cover crops on nitrogen leaching, net greenhouse gas balance and crop productivity. Global change biology, 25(8), 2530–2543.

2 Albornoz, F. E., Hayes, P. E., Orchard, S., Clode, P. L., Nazeri, N. K., Standish, R. J., … & Ryan, M. H. (2020). First cryo-scanning electron microscopy images and X-ray microanalyses of mucoromycotinian fine root endophytes in vascular plants. Frontiers in Microbiology, 11, 2018.

3 Albornoz, F. E., Orchard, S., Standish, R. J., Dickie, I. A., Bending, G. D., Hilton, S., … & Ryan, M. H. (2021). Evidence for niche differentiation in the environmental responses of co-occurring Mucoromycotinian fine root endophytes and Glomeromycotinian arbuscular mycorrhizal fungi. Microbial Ecology, 81, 864–873.

4 Altieri, M. A., Nicholls, C. I., Henao, A., & Lana, M. A. (2015). Agroecology and the design of climate change-resilient farming systems. Agronomy for sustainable development, 35(3), 869–890.

5 Arines, J., Vilarino, A., & Sainz, M. (1988). ‘Fine’and ‘coarse’mycorrhizal fungi on red clover plants in acid soils: Root colonization and plant responses. Plant and soil, 111, 135–145.

6 Bonfante, P., & Venice, F. (2020). Mucoromycota: going to the roots of plant-interacting fungi. Fungal Biology Reviews, 34(2), 100–113.

7 Cole, J., Raguideau, S., Abbaszadeh-Dahaji, P., Hilton, S., Muscatt, G., Mushinski, R. M., … & Bending, G. D. (2024). Comparative genomic analysis of a metagenome-assembled genome reveals distinctive symbiotic traits in a Mucoromycotina fine root endophyte arbuscular mycorrhizal fungus. bioRxiv, 2024–11.

8 Crush, J. R. (1975). Occurrence of endomycorrhizas in soils of the Mackenzie Basin, Canterbury, New Zealand. New Zealand journal of agricultural research, 18(4), 361–364.

9 Douds Jr, D. D., & Millner, P. D. (1999). Biodiversity of arbuscular mycorrhizal fungi in agroecosystems. Agriculture, ecosystems & environment, 74(1-3), 77–93.

10 Field, K. J., Bidartondo, M. I., Rimington, W. R., Hoysted, G. A., Beerling, D., Cameron, D. D., … & Pressel, S. (2019). Functional complementarity of ancient plant–fungal mutualisms: contrasting nitrogen, phosphorus and carbon exchanges between Mucoromycotina and Glomeromycotina fungal symbionts of liverworts. New Phytologist, 223(2), 908–921.

11 Frey, S. D. (2019). Mycorrhizal fungi as mediators of soil organic matter dynamics. Annual review of ecology, evolution, and systematics, 50(1), 237–259.

12 Gange, A. C. (2007). Insect-mycorrhizal interactions: patterns, processes, and consequences. Ecological communities: plant mediation in indirect interaction webs, 124–143.

13 Giovannetti, M., & Mosse, B. (1980). An evaluation of techniques for measuring vesicular arbuscular mycorrhizal infection in roots. New phytologist, 489–500.

14 Howard, N. O., Williams, A., Durant, E., Pressel, S., Daniell, T. J., & Field, K. J. (2024). Preferential nitrogen and carbon exchange dynamics in Mucoromycotina “fine root endophyte”-plant symbiosis. Current Biology.

15 Hoysted, G. A., Field, K. J., Sinanaj, B., Bell, C. A., Bidartondo, M. I., & Pressel, S. (2023). Direct nitrogen, phosphorus and carbon exchanges between Mucoromycotina ‘fine root endophyte’fungi and a flowering plant in novel monoxenic cultures. New Phytologist, 238(1), 70–79.

16 Ivanov, S., & Harrison, M. J. (2014). A set of fluorescent protein-based markers expressed from constitutive and arbuscular mycorrhiza-inducible promoters to label organelles, membranes and cytoskeletal elements in Medicago truncatula. The Plant Journal, 80(6), 1151–1163.

17 Jeffery, R. P., Simpson, R. J., Lambers, H., Orchard, S., Kidd, D. R., Haling, R. E., & Ryan, M. H. (2018). Contrasting communities of arbuscule-forming root symbionts change external critical phosphorus requirements of some annual pasture legumes. Applied Soil Ecology, 126, 88–97.

18 Karandashov, V., & Bucher, M. (2005). Symbiotic phosphate transport in arbuscular mycorrhizas. Trends in plant science, 10(1), 22–29.

19 Klironomos, J. N., & Hart, M. M. (2002). Colonization of roots by arbuscular mycorrhizal fungi using different sources of inoculum. Mycorrhiza, 12, 181–184.

20 Kobae, Y., Tamura, Y., Takai, S., Banba, M., & Hata, S. (2010). Localized expression of arbuscular mycorrhiza-inducible ammonium transporters in soybean. Plant and Cell Physiology, 51(9), 1411–1415.

21 Kowal, J., Arrigoni, E., Serra, J., & Bidartondo, M. (2020). Prevalence and phenology of fine root endophyte colonization across populations of Lycopodiella inundata. Mycorrhiza, 30, 577–587.

22 Lutz, S., Mikryukov, V., Labouyrie, M., Bahram, M., Jones, A., Panagos, P., Delgado-Baquerizo, M., Maestre, F. T., Orgiazzi, A., Tedersoo, L., & van der Heijden. M.G.A. (2025). Global richness of arbuscular mycorrhizal fungi. Fungal Ecology, 74 101407, ISSN 1754-5048, 10.1016/j.funeco.2024.101407.

23 Orchard, S., Hilton, S., Bending, G. D., Dickie, I. A., Standish, R. J., Gleeson, D. B., … & Ryan, M. H. (2017a). Fine endophytes (Glomus tenue) are related to Mucoromycotina, not Glomeromycota. New Phytologist, 213(2), 481–486.

24 Orchard, S., Standish, R. J., Dickie, I. A., Renton, M., Walker, C., Moot, D., & Ryan, M. H. (2017b). Fine root endophytes under scrutiny: a review of the literature on arbuscule-producing fungi recently suggested to belong to the Mucoromycotina. Mycorrhiza, 27, 619–638.

25 Orchard, S., Standish, R. J., Nicol, D., Dickie, I. A., & Ryan, M. H. (2017c). Sample storage conditions alter colonisation structures of arbuscular mycorrhizal fungi and, particularly, fine root endophyte. Plant and Soil, 412, 35–42.

26 Orchard, S., Standish, R. J., Nicol, D., Gupta, V. V. S. R., & Ryan, M. H. (2016). The response of fine root endophyte (Glomus tenue) to waterlogging is dependent on host plant species and soil type. Plant and Soil, 403, 305–315.

27 Pearson, J. N., Smith, S. E., & Smith, F. A. (1991). Effect of photon irradiance on the development and activity of VA mycorrhizal infection in Allium porrum. Mycological Research, 95(6), 741–746.

28 Powell, J. R., & Rillig, M. C. (2018). Biodiversity of arbuscular mycorrhizal fungi and ecosystem function. New phytologist, 220(4), 1059–1075.

29 Prout, J. N., Williams, A., Wanke, A., Schornack, S., Ton, J., & Field, K. J. (2024). Mucoromycotina ‘fine root endophytes’: a new molecular model for plant–fungal mutualisms?. Trends in Plant Science, 29(6), 650–661.

30 Rimington, W. R., Pressel, S., Duckett, J. G., & Bidartondo, M. I. (2015). Fungal associations of basal vascular plants: reopening a closed book?. New Phytologist, 205(4), 1394–1398.

31 Ruess, R. W., Hendrick, R. L., Burton, A. J., Pregitzer, K. S., Sveinbjornssön, B., Allen, M. F., & Maurer, G. E. (2003). Coupling fine root dynamics with ecosystem carbon cycling in black spruce forests of interior Alaska. Ecological Monographs, 73(4), 643–662.

32 Seeliger, M., Hilton, S., Muscatt, G., Walker, C., Bass, D., Albornoz, F., … & Bending, G. D. (2024). New fungal primers reveal the diversity of Mucoromycotinian arbuscular mycorrhizal fungi and their response to nitrogen application. Environmental microbiome, 19(1), 71.

33 Serrano, K., Bezrutczyk, M., Goudeau, D., Dao, T., O’Malley, R., Malmstrom, R. R., … & Cole, B. (2024). Spatial co-transcriptomics reveals discrete stages of the arbuscular mycorrhizal symbiosis. Nature Plants, 1–16.

34 Sinanaj, B., Pressel, S., Bidartondo, M. I., & Field, K. J. (2024). Fungal symbiont diversity drives growth of Holcus lanatus depending on soil nutrient availability. Functional Ecology, 38(4), 984–997.

35 Thippayarugs, S., Bansal, M., & Abbott, L. K. (1999). Morphology and infectivity of fine endophyte in a Mediterranean environment. Mycological Research, 103(11), 1369–1379.

36 Turner Ii, B. L., & Meyer, W. B. (1991). Land use and land cover in global environmental change: considerations for study. International Social Science Journal, 43(129).

37 Walker, C., Gollotte, A., & Redecker, D. (2018). A new genus, Planticonsortium (Mucoromycotina), and new combination (P. tenue), for the fine root endophyte, Glomus tenue (basionym Rhizophagus tenuis). Mycorrhiza, 28(3), 213–219.

