## Supplementary Figure 1 for "Fine root endophytes forming winter mycorrhiza"

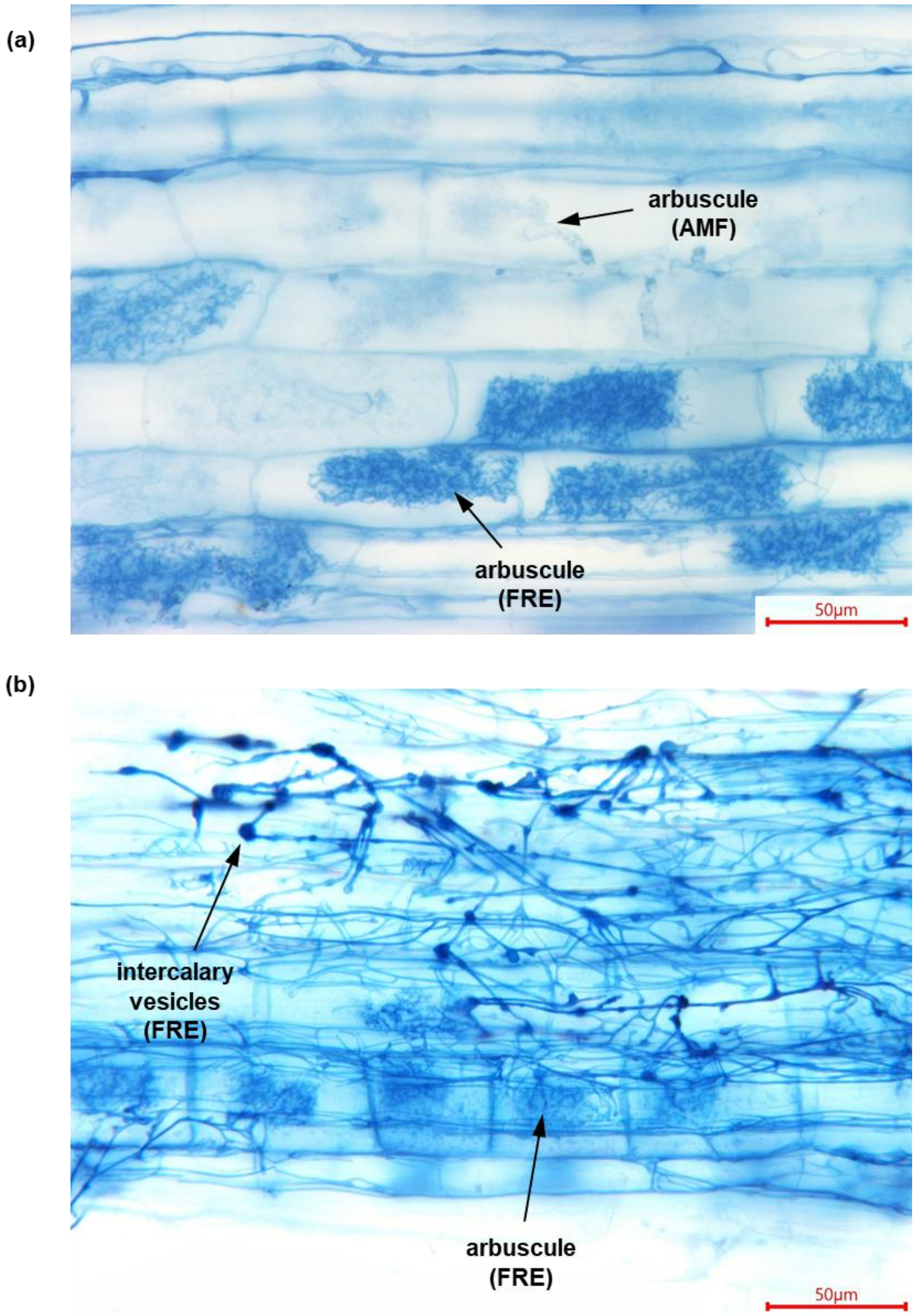


**Supplementary Figure 1, Fine root endophytes morphology**

**(a)** *Erigeron annuus* (L) mycorrhizal root stained with trypan blue; Fine root endophytes (FRE) is darkly stained, whereas co-colonizing arbuscular mycorrhizal fungi (AMF) is lightly stained. Roots sampled in November 2024 **(b)** *Lactuca sativa* (L) mycorrhiza stained with trypan blue; Numerous intercalary vesicles, characteristic of FRE, are observed on the thin intraradical hyphae. Roots sampled in January 2025
